# k2v: A Containerized Workflow for Creating VCF Files from Kintelligence Targeted Sequencing Data

**DOI:** 10.1101/2022.11.21.517402

**Authors:** Stephen D. Turner, Michelle A. Peck

## Abstract

The ForenSeq Kintelligence kit developed by Verogen is a targeted Illumina sequencing assay that genotypes 10,230 single nucleotide polymorphisms designed for forensic genetic genealogy, forensic DNA phenotyping, and ancestry inference. We developed `k2v`, a containerized workflow for creating standard specification-compliant variant call format (VCF) files from the custom output data produced by the Kintelligence Universal Analysis Software. VCF files produced with `k2v` enable the use of many pre-existing, widely used, community-developed tools for manipulating and analyzing genetic data in the standard VCF format. Here we describe the `k2v` implementation, demonstrate its usage, and use the VCF produced by `k2v` to demonstrate downstream analyses that can easily be performed with pre-existing tools using VCF data as input: concordance analysis, ancestry inference, and relationship estimation. `k2v` is distributed as a Docker container available on Docker Hub. Documentation and source code for `k2v` is freely available under the GNU Public License (GPL-3.0) at https://github.com/signaturescience/k2v.

## 1 Introduction

Kinship analysis using single nucleotide polymorphisms (SNPs) is a cornerstone of forensic genetic genealogy (FGG) analysis (Greytak, Moore, and Armentrout 2019; Kling and Tillmar 2019; Glynn 2022). Originally used by hobbyist and professional genealogists, databases populated with direct-to-consumer DNA tests such as GEDmatch and Family Tree DNA are now widely used by law enforcement agencies to generate investigative leads on unknown crime scene samples or unidentified human remains by finding distant relatives of the unknown DNA sample. The technique has gained enough traction in the community to prompt the Federal Bureau of Investigation (FBI) to call for self-regulation and development of guidelines for practicing responsible genetic genealogy (Callaghan 2019).

FGG routinely relied on genome-wide SNP data using whole genome sequencing or microarrays (Russell et al. 2022; de Vries et al. 2022; Davawala et al. 2022; Tillmar et al. 2020). However, targeted sequencing assays have recently been developed that leverage a sparse set of SNPs for kinship analysis, including the community-developed FORCE panel (Tillmar et al. 2021) and the commercially available ForenSeq Kintelligence kit (Verogen, Inc., San Diego CA), which we internally validated (Peck et al. 2022).

The Kintelligence kit provides genotypes on 10,230 SNPs, of which 9,867 are kinship-informative SNPs used for SNP-based kinship analysis across populations, while the remaining 363 are used to inform biogeographical ancestry, identity, hair and eye color, and biological sex (Snedecor et al. 2022). The Kintelligence kit integrates data analysis in a browser-based application, and also produces Excel and text file outputs containing genotype data in a custom format.

The Variant Call Format (VCF) is the *de facto* standard file format for storing genetic variant data that has been used by genetics and bioinformatics community for over a decade (P. Danecek et al. 2011). A vast ecosystem of widely used, community-developed, and community-supported tools exist for managing and analyzing variant data in VCF files including tools for general data manipulation and analysis (Petr Danecek et al. 2021), and tools specifically designed for forensic genetics-focused analysis (Turner et al. 2022; V. P. Nagraj et al. 2022b).

In this paper we describe a new tool, `k2v`, for converting Kintelligence data into a standard specification-compliant VCF, which enables the use of a vast pre-existing infrastructure for genetic data manipulation and analysis. The `k2v` workflow is implemented as a Docker container, and can be run on any system where Docker or Singularity is available. We provide a demonstration of the `k2v` workflow and several example forensic genetics-relevant analyses that can easily be accomplished using standard tools that use VCF data as input.

## 2 Implementation

`k2v` is implemented as a Docker container and is distributed to run directly from a command line interface. The `k2v` Docker image has bcftools (Petr Danecek et al. 2021), htslib (Bonfield et al. 2021), and R (R Core Team 2017) installed on a lightweight Alpine Linux base image. Running the `k2v` container instantiates a script inside the container that takes as input the custom `*.SnpResult.txt` file that is produced by Kintelligence, joins that data to a table of reference and alternate alleles obtained by extracting Kintelligence sites from the GnomAD site VCF (Karczewski et al. 2020) to create an intermediate tabular file, which is then converted to VCF using bcftools, and written out to the host filesystem after being compressed with bgzip and indexed with tabix. Documentation and implementation details are available in the project repository at https://github.com/signaturescience/k2v.

## 3 Use cases

### 3.1 Validation data analysis

With only the custom format text file produced by the Kintelligence platform, custom analysis workflow development would be required for basic data analysis tasks such as call rate, concordance, and heterozygosity analysis that may be used in a validation study. Instead, we can convert the Kintelligence data into VCF then use pre-existing tools for data analysis to streamline analysis.

In this example we convert Kintelligence data generated on the NIST Standard Reference Material sample HG002 (NA24385) to VCF using `k2v`. We first use bcftools (Petr Danecek et al. 2021) to calculate simple statistics on the VCF. We then use the previously published `nrc` tool (V. P. Nagraj et al. 2022b) to get a detailed analysis of how the Kintelligence genotype data differs from the benchmark callset from NIST/Genome-in-a-Bottle. Additional details on `k2v` usage and getting example data can be found on the `k2v` GitHub README. This analyses required less than one second of compute time. Table 1 shows the count of the number of substitutions and variant types in sample HG002. Table 2 shows a subset of results from running the `nrc` tool on the VCF produced by `k2v`, comparing the Kintelligence VCF to the GIAB high-confidence call set.

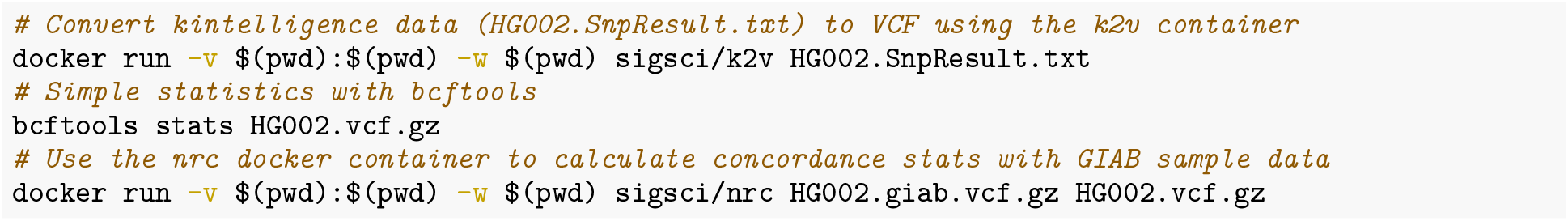

**Table 1:** Count of the number of substitutions and variant types in sample HG002.

| Substitution / variant type | Count |
| --- | --- |
| A>C | 288 |
| A>G | 1,439 |
| A>T | 4 |
| C>A | 289 |
| C>G | 4 |
| C>T | 1,614 |
| G>A | 1,515 |
| G>C | 2 |
| G>T | 302 |
| T>A | 1 |
| T>C | 1,424 |
| T>G | 307 |
| Homozygous REF | 2,897 |
| Homozygous ALT | 2,590 |
| Heterozygous | 4,599 |
| Transitions | 5,992 |
| Transversions | 1,197 |
| Missing | 55 |

**Table 2:** Subset of output from the `nrc` tool comparing the Kintelligence data converted to VCF using `k2v` against the high-confidence GIAB callset from HG002. The table shows the total number of biallelic SNPs compared between the two samples, discordance and concordance statistics, non-reference discordance (NRD), non-reference concordance (NRC), and the full array of each type of genotype match and mismatch. The vector of match/mismatch counts can be interpreted as follows: RRRR = number of genotype matches where the GIAB and Kintelligence calls are both homozygous for the reference allele; RRRA = number of genotype mismatches where the GIAB call is homozygous reference (Ref/Ref), but the Kintelligence VCF is heterozygous (Ref/Alt); RARA = number of genotype matches where both the GIAB callset and the Kintelligence VCF are both heterozygous (Ref/Alt); etc. See (V. P. Nagraj et al. 2022b) for definitions of NRD and NRC. Note that the total number of SNPs compared is fewer than the number of SNPs assayed by Kintelligence – this is because the GIAB call set only includes autosomes, and the high-confidence regions do not cover all the Kintelligence-assayed sites.

| Metric | Value |
| --- | --- |
| Total | 9,802 |
| Discordance | 0.00867 |
| Concordance | 0.991 |
| NRD | 0.0121 |
| NRC | 0.988 |
| RRRR | 2,779 |
| RRRA | 2 |
| RRAA | 0 |
| RARR | 40 |
| RARA | 4,511 |
| RAAA | 42 |
| AARR | 0 |
| AARA | 1 |
| AAAA | 2,427 |

### 3.2 Ancestry inference

The use of principal components analysis (PCA) for studying genetic variation data was introduced over 40 years ago (Menozzi, Piazza, and Cavalli-Sforza 1978). Since its introduction PCA has been widely used in population and medical genetics for biogeographic ancestry analysis (Patterson, Price, and Reich 2006; Novembre et al. 2008; Li et al. 2008; Wellcome Trust Case Control Consortium 2007; Tian et al. 2008; Price et al. 2008; Shriver and Kittles 2004), and has recently gained in the forensic genomics community for statistical inference of an unknown DNA donor’s ancestry (Phillips 2015).

There are several existing tools for conducting PCA with genetic variant data including EIGENSTRAT (Price et al. 2006) and PLINK (Purcell et al. 2007), both widely used and well-supported by the genetic epidemiology community. The Ancestry and Kinship Tools (AKT) developed by Illumina is a permissively licensed open-source implementation of PCA (and other analyses relevant for forensic genomics), which uses the htslib API allowing it to work directly with VCF files.

In this example we use AKT to perform PCA on the HG002 VCF produced by Kintelligence against the 2,504 individuals in the 1000 Genomes Project phase 3 release data. In this example we start with the 1000 Genomes data on the kintelligence sites, merge in the HG002 VCF produced by `k2v`, then run PCA using AKT. We read in the output into R, annotate the 1000 Genomes samples with continental ancestry (Turner 2022). This analysis required less than 10 seconds of compute time. Figure 1 shows a PCA biplot showing the first two principal components, with HG002 highlighted against the background of 1000 Genomes samples.

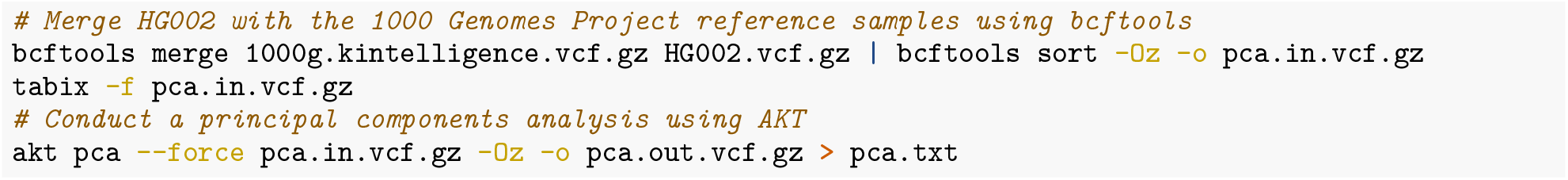

**Figure 1:**
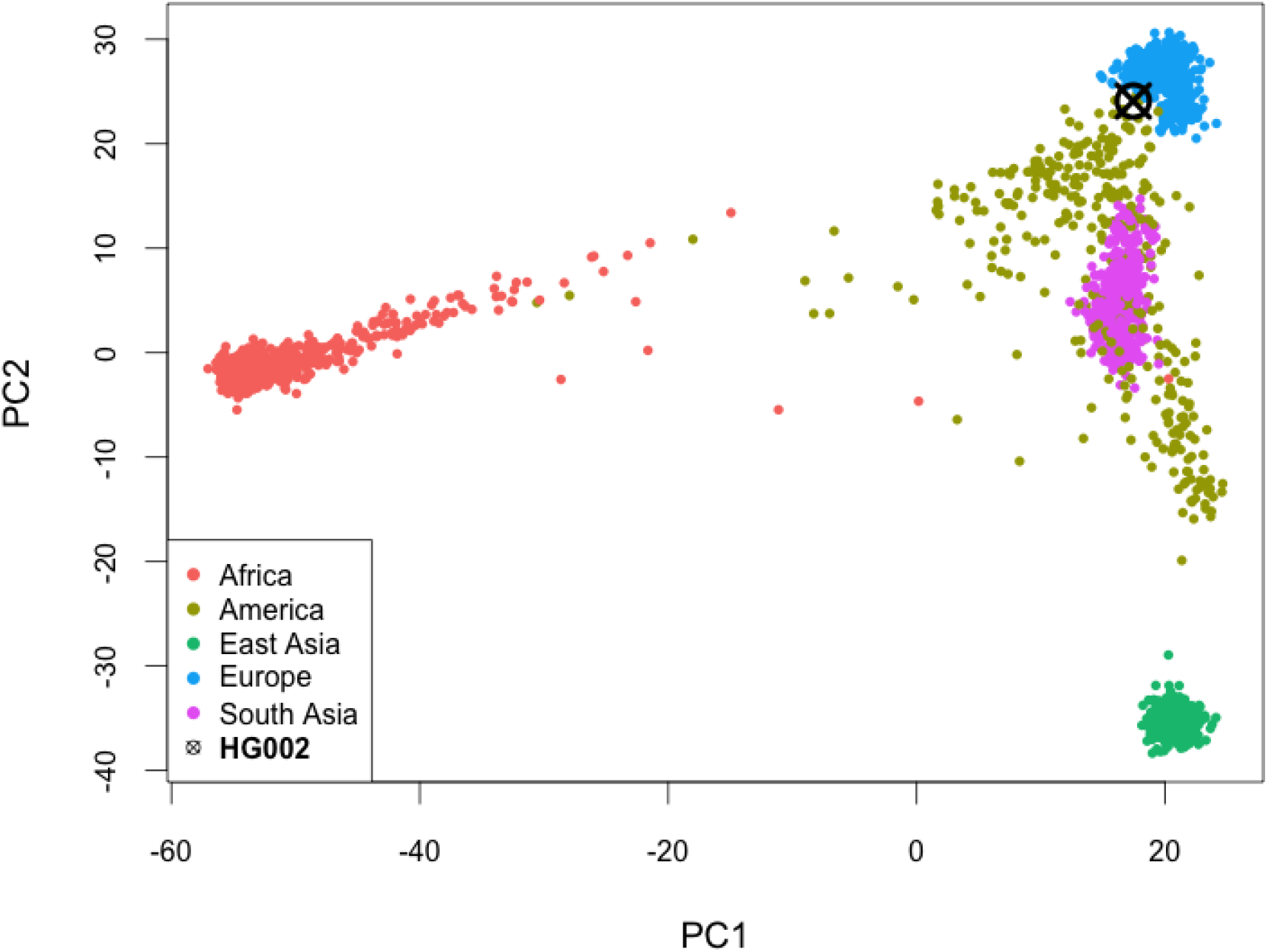
Principal components analysis (PCA) biplot, showing the first two principal components highlighting the HG002 VCF produced by `k2v` against the 2,504 samples in the 1000 Genomes Project phase 3 release data.

### 3.3 Relationship estimation

Relationship inference methods using genome-wide SNP data typically fall into two broad classes of methods: measures that directly infer relatedness genome-wide, and identity by descent (IBD) segment approaches (Ramstetter et al. 2017). These approaches to relationship inference have been benchmarked using data generated in ideal conditions (Ramstetter et al. 2017), and more recently assessed using low-quality microarray data on forensic samples (Turner et al. 2022) or with low-coverage whole genome sequencing data followed by imputation (V. P. Nagraj et al. 2022a).

In this example, we use the KING-robust estimator (Manichaikul et al. 2010) implemented in the PLINK 2 software (Chang et al. 2015) to estimate relatedness between two Ashkenazi Jewish individuals from the Personal Genome Project: HG002 (NA24385; the son), and HG004 (NA24143, the mother), using VCFs produced by `k2v` from Kintelligence data. This analysis required less than 50 milliseconds of compute time.

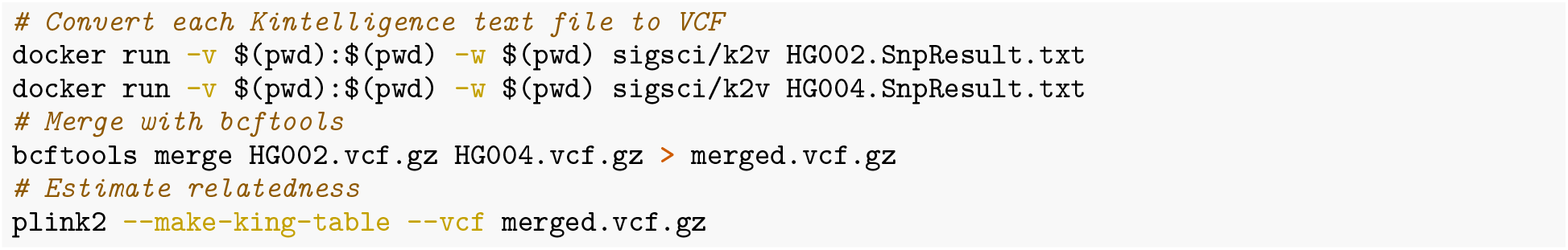

The kinship coefficient between these two samples as inferred using the KING-robust estimator is *φ* = 0.2384, which clearly indicates that this sample pair is a first-degree relationship. The proportion of loci where zero alleles are shared identical by state is *κ*_0_ = 0.0039. Because parent-child offspring are expected to share *κ*_0,1,2_ = (0, 1, 0) while full siblings are expected to share 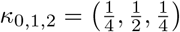, we correctly infer that HG002 and HG004 are a parent-offspring pair.

## 4 Conclusion

To use data from a sequencing or genotype assay that produces variant calls in a custom format requires either (a) custom development of new analysis tools for each downstream analysis task, or (b) development of a single tool to convert data to a common format for which tools already exist. `k2v` takes the latter approach, enabling the rapid conversion of Kintelligence variant data in a custom format to a standard specification-compliant VCF file. Once data exists in a VCF, nearly any downstream analyses and manipulation tasks can readily be achieved with existing infrastructure. `k2v` is currently limited to SNPs only, and cannot capture the one insertion that Kintelligence targets (rs796296176).

`k2v` is freely available under the GNU Public License (GPL) v3 at https://github.com/signaturescience/k2v with further documentation on the repository README. A pre-built Docker image is available on Docker hub and can be installed via `docker pull sigsci/k2v`.

## Author contributions

**Stephen Turner**: Conceptualization, Methodology, Software, Validation, Investigation, Resources, Data Curation, Writing – Original Draft, Writing – Review & Editing, Visualization, Project Administration.

**Michelle Peck**: Resources, Data Curation, Writing – Review & Editing.

## Author disclosure statement

The authors have no competing financial interests to disclose. Any mention of commercial products was done for scientific transparency and should not be viewed as an endorsement of the product or manufacturer.

## Acknowledgements

The authors thank Erin Gorden, Christina Neal, Dr. Carmen Reedy, and Dr. Alex Koeppel for helpful discussions on the utility of this tool.

## Funding

This research received no external funding.

## Notes

https://github.com/signaturescience/k2v

https://hub.docker.com/r/sigsci/k2v

### Competing Interest Statement

The authors have declared no competing interest.

